# Dissection of the contribution of regulation mode to the properties of feedforward and feedback gene regulatory motifs

**DOI:** 10.1101/065698

**Authors:** Bharat Ravi Iyengar, Beena Pillai, K.V. Venkatesh, Chetan J. Gadgil

**Author notes:** (Chetan J. Gadgil).

## Abstract

We present a framework enabling dissection of the effects of motif structure (feedback or feedforward), nature of the controller (mRNA or protein), and regulation mode (transcriptional, post-transcriptional or translational) on the response to a step change in the input. We have used a common model framework for gene expression where both motif structures have an activating input and repressing regulator, with the same set of parameters to enable comparison of the responses. We studied the global sensitivity of the system properties such as steady-state gain, overshoot, peak time, and peak duration, to parameters. We find that, in all motifs, overshoot correlated negatively whereas peak duration varied concavely, with peak time. Differences in other system properties were found to be mainly dependent on the the nature of the regulator, than the motif structure. Protein mediated motifs showed a higher degree of adaptation; feedforward motifs exhibited perfect adaptation. RNA mediated motifs had a mild regulatory effect; they also exhibited lower peaking tendency and mean overshoot. Protein mediated feedforward motifs showed higher overshoot and lower peak time compared to corresponding feedback motifs.

## 1 Introduction

Gene regulation is one of the central processes that dictates the behaviour of a cell and its responsiveness to the environment. Since gene expression is a multi-step process, different modes of regulation can affect the steady state of a gene product by either controlling the formation rate or the degradation rate of the transcripts or the proteins^[1]^. The controller employed by the regulatory mechanism could be a protein, or transcripts such as different classes of non-coding RNAs^[2, 3]^.

The interaction network of all the genes in the genome is called the gene regulatory network (GRN). Despite its complexity, the GRN has a higher representation of certain subnetworks that have a distinct pattern of connections known as network motifs^[4]^. Feedback and feedforward loops are two of the most commonly occuring network motifs in the GRN^[5, 6]^.

In a feedback loop, a gene regulates its own activity and depending on whether the regulation is repressive or activating, the motif is either called a negative or a positive feedback loop, respectively. Feedbacks can be direct, in which the output influences its own activity, or indirect, in which it acts via an intermediary (controller). The same regulatory action, for example a negative effect of the controller on the output, can be implemented via several functionally equivalent motifs.

In feedforward loops, the input regulates the output simultaneously via two regulatory paths. If the effects of the two paths on the regulated gene are the same then the motif is called a coherent feedforward motif and otherwise, an incoherent feedforward motif. For example the level of the output would be governed directly by the input signal (input) and indirectly via the controller.

In this study we focus our analyses, for both feedforward and feedback motifs, on those cases in which the action of the input is non-negative and that of the controller is non-positive. Under most operating conditions, the input is expected to cause an increase (or no change) in the output signal for an unregulated system. Similarly, an increase in the magnitude of the controller action is expected to result in a decrease (or at the most no change) of the output signal for a constant level of the input signal. In particular, we have modelled and compared the indirect negative feedback loop (NFBL) and type-1 incoherent feedforward loop (I1-FFL) network motifs (Fig. 1), in both of which there are three nodes and the controller action is negative.

**Fig. 1:**
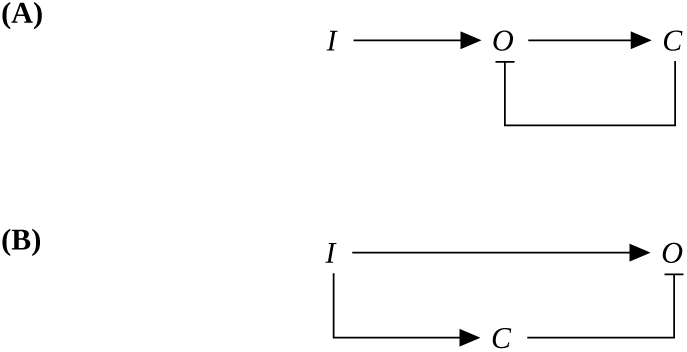
The (**A**) Indirect Negative Feedback and (**B**) Type-1 Incoherent Feedforward loop, with three nodes. *I* refers to Input, *O* refers to output and *C* refers to Controller. The input has a positive effect and the controller has a negative effect on the output in both the motifs.

The I1-FFL motif is known to generate a dynamic “pulse” in the output gene expression under certain conditions. The shape and dynamics of the pulse depends on various parameters such as the strength of activation or repression, and other kinetic rate constants. If the strength of the repression is strong enough then the output expression may return back to the pre-activation levels; this phenomenon is called adaptation^[8]^.

NFBLs are also implicated in adaptation and it has been shown using a mathematical model for a bacterial chemotaxis system that negative feedback via integral control can lead to a perfect adaptation^[14]^. Ma et al. have extensively studied different variants of feedforward and feedback motifs that lead to adaptation^[12]^. In another important study by Rosenfeld et al., it was shown that negative autoregulatory motif has a shortened response time, compared to the unregulated system, given the constraint that both the systems have the same steady state level of the output^[15]^. However, these models have not explicitly studied the effect of mode of regulation or nature of the controller, on the dynamics and steady state properties of the motif.

Robustness is a property of a system by virtue of which a system can maintain its output, for a given input, despite changes in intrinsic parameters. Most common techniques of assessing robustness using mathematical models include a local sensitivity analysis at the steady state and parameter variation to study global effects. In a study on the robustness of different feedforward motifs by Wang et al., it was concluded that I1-FFL is one of the most robust feedforward motifs^[10]^. Integral feedback control in the bacterial chemotaxis model was also found to exhibit some robustness^[14]^. Iadevaia et al., using mass action kinetics based models, have studied the robustness of the output gene expression for different types feedback, feedforward and mixed motifs in cell signalling, to variations in parameters^[13]^. A summary of these different models and analyses is presented in Tab. 1.

**Tab. 1:** Summary of previous models on feedback and/or feedforward motifs.

| Reference | Motifs studied | Metrics used | Mode of regulation | Nature of regulator | Parameter Sampling |
| --- | --- | --- | --- | --- | --- |
| Mangan & Alon<sup>[7]</sup> | FFLs | Response Time | ✗ | ✗ | ✗ |
| Sontag<sup>[8]</sup> | I1-FFL | Adaptation | ✗ | ✗ | ✗ |
| Goentoro et al.<sup>[9]</sup> | I1-FFL | Adaptation | ✗ | ✗ | ✓<br>(selected) |
| Wang et al.<sup>[10]</sup> | FFLs | Input-Output sensitivity | ✗ | ✗ | ✓<br>(selected) |
| Duk et al.<sup>[11]</sup> | FFLs | ✗ | ✓ | ✓ | ✓<br>(selected) |
| Ma et al.<sup>[12]</sup> | FFLs, FBLs | Adaptation | ✗ | ✗ | ✓<br>(method not described) |
| Iadevaia et al.<sup>[13]</sup> | FFLs, FBLs, mixed | Robustness | ✗ | ✗ | ✓<br>(random) |

Since the role of RNA mediated regulation in cellular processes is becoming increasingly evident, it is essential to delineate their roles from strictly protein mediated regulation, as a part of these motifs. A significant number of miRNA mediated gene regulatory pathways are also enriched with feedback and feedforward motifs^[16]^. Duk et al. have extensively studied the dynamics of different types miRNA mediated FFLs with different underlying mechanistic models^[11]^. However, they have limited their analyses to just FFLs and a comparison with FBLs with similar regulatory mechanism is not possible with their model.

Though all these models and analyses have provided fundamental insights about the properties of feedback and feedforward motifs, a systematic comparison of motif properties along with the effect of parameter variation has not been done using a common mechanistic model. Some of these studies have mentioned and analysed both feedback and feedforward motifs but have not carried out an explicit comparison between them. Also, many of these studies have modelled a generic network and have not attributed any biological mechanism. To our knowledge, there is no study that uses a mechanism-based model to simultaneously evaluate the contributions of motif structure, mode of regulation and the nature of controller, on the dynamics and steady state response of the output. Therefore, we have developed a framework to simultaneously compare different properties of protein or RNA mediated genetic feedforward (I1-FFL) and feedback (NFBL) motifs using defined metrics. All these motifs share a basic set of parameters which was chosen from realistic estimates obtained from experimental data reported in the available literature. The sensitivities of the different properties to parameter changes have been studied by random multivariate sampling of all the parameters in a defined range which is more informative than variation of individual parameters. Our model reveals some previously unexplored features of feedback and feedforward motifs while corroborating some of the known properties. We find that while some properties are dependent on motif structure, others are strongly dependent on the nature of regulator. For every result, we also assessed the effect of the choice of parameters, in particular the protein degradation rate and therefore abundance, which is known to vary over multiple orders of magnitude^[17]^.

## 2 Methods

### 2.1 Mathematical description of the response metrics

Mathematical models were developed for feedforward (FFL) motifs and feedback motifs (FBL) each regulated by either a protein or an RNA at transcriptional, post-transcriptional or translational levels. These twelve combinations are abbreviated using a six letter code as indicated in Tab. 2.

**Tab. 2:**
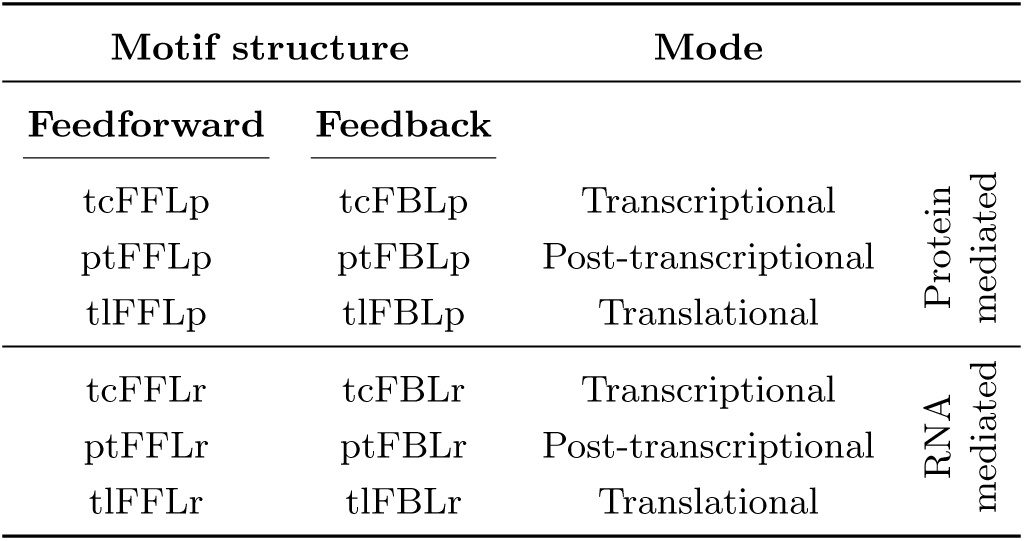
Abbreviations for different motifs. First two letters of the abbreviation indicate the mode of regulation, the next three (in caps) denote the type of motif (feedback/feedforward) and the last letter represents the nature of the controller (protein/RNA).

| Motif structure |  | Mode |  |
| --- | --- | --- | --- |
| Feedforward | Feedback |  |  |
| tcFFLp | tcFBLp | Transcriptional | Protein mediated |
| ptFFLp | ptFBLp | Post-transcriptional |  |
| tlFFLp | tlFBLp | Translational |  |
| tcFFLr | tcFBLr | Transcriptional | RNA mediated |
| ptFFLr | ptFBLr | Post-transcriptional |  |
| tlFFLr | tlFBLr | Translational |  |

All motifs (Fig. 2) contain four components – output RNA (*r*_*out*_), controller RNA (*r*_*ctrl*_), output protein (*p*_*out*_) and controller protein (*p*_*ctrl*_). Since unregulated motif does not have any effect of the controller, *r*_*ctrl*_ and *p*_*ctrl*_ are decoupled from the other components. Likewise, in RNA mediated motifs, there is no role of the *p*_*ctrl*_ other than as a marker for *r*_*ctrl*_ levels and is therefore not considered in further analyses.

**Fig. 2:**
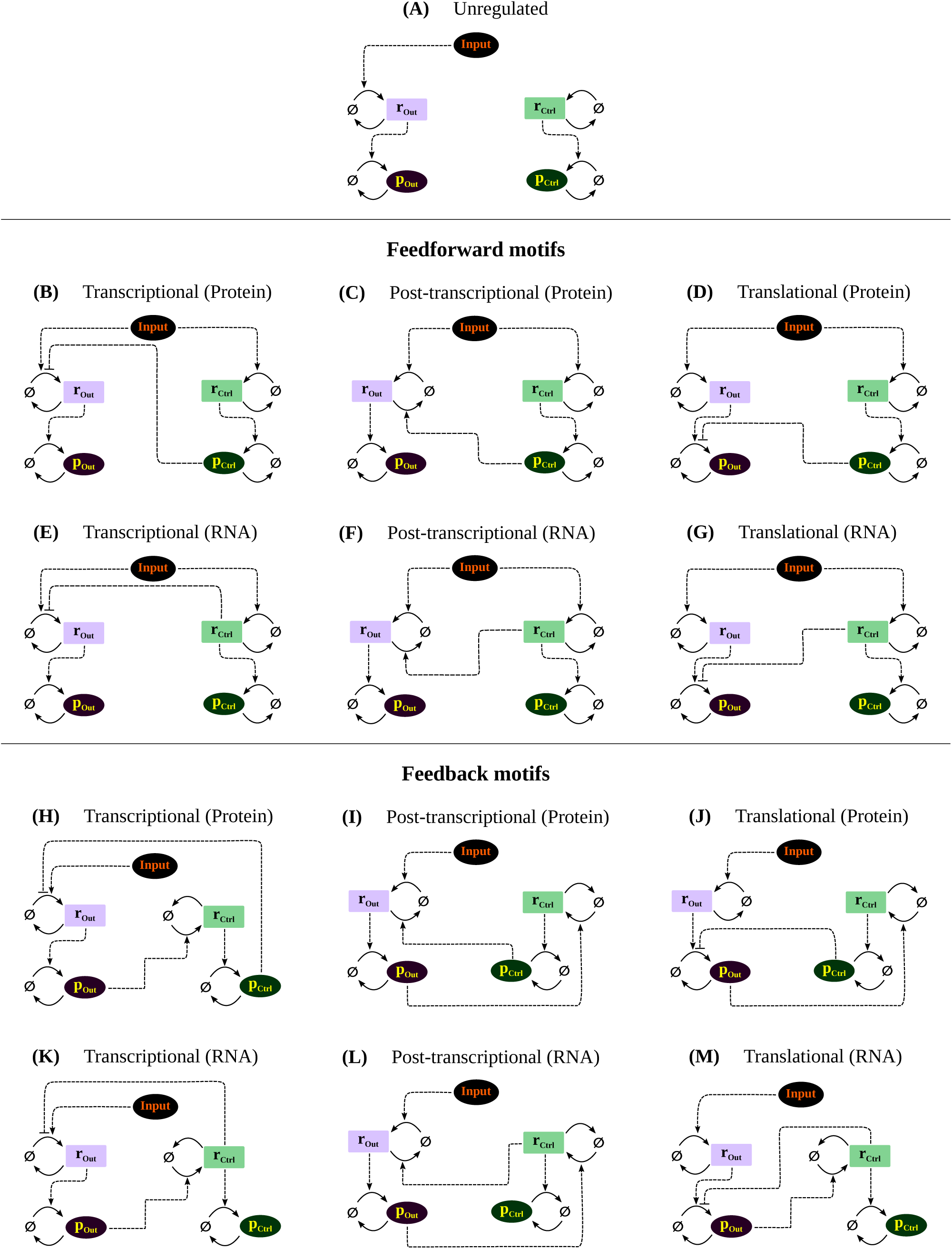
Motifs: (**A**) Unregulated. Protein (**B-D**) and RNA mediated (**E-G**) feedforward motifs. Protein (**H-J**) and RNA mediated (**K-M**) feedback motifs. Output mRNA, Output protein, Controller mRNA and Controller protein are denoted as *r*_*out*_(lilac box), *p*_*out*_ (purple oval), *r*_*ctrl*_ (light green box) and *p*_*ctrl*_ (deep green oval), respectively. Ø denotes the cellular pool of nucleotides and amino acids, which is assumed to be infinite.

Transcription of *r*_*out*_ is activated by an input which in turn leads to production of *p*_*out*_. In transcriptional regulation, the controller, either RNA or protein, inhibits the transcription of *r*_*out*_ whereas in translational regulation the controller inhibits the production of *p*_*out*_. Post-transcriptional motifs are modelled such that they promote degradation of *r*_*out*_. In feedforward motifs the transcription of controller RNA is activated by the input whereas in feedback motifs it is activated by the output protein.

The responses of these motifs to a step change in the input were studied. At time, t=0, the system was initialized at the steady state for a given value of input (initial state). The input was stepped up and the dynamics of the response was monitored. The motifs were compared using different metrics (Fig. 3) which are defined as follows.

**Fig. 3:**
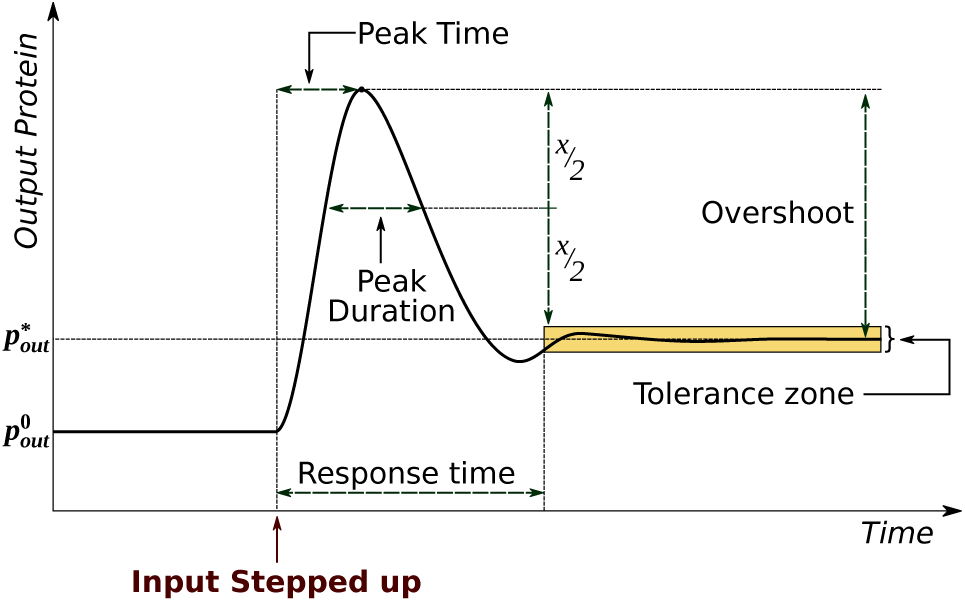
Illustration of the metrics measured after dynamic simulation of different motifs. These metrics were used for the comparison of these motifs. Steady state gain is not shown in this picture.

Steady state gain is defined as the ratio of the change in the steady state level and the initial steady state level of the output protein^[18]^.

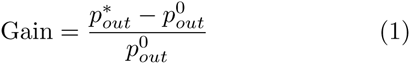

Here, 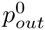 and 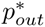 denote the initial and final steady state values of *p*_*out*_, respectively.

Response time (*t*_resp_), also called settling time, is the time taken by the output protein to reach a defined tolerance zone of ±1% of the steady state^[19, 18]^.

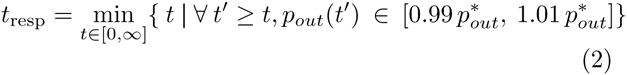

Overshoot is defined as the difference between the maximum peak height and the steady state level of the output protein. This value was normalized to the steady state to obtain a per-unit overshoot^[19]^.

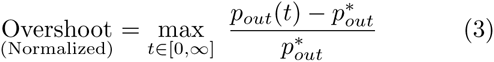

Peak time is defined as the time taken to reach the maximum peak value^[18]^. For the analyses peak time was normalized by the response time.

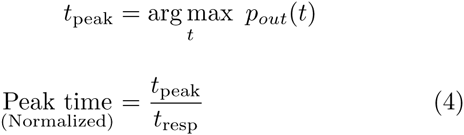

Peak duration represents the width of the peak and is defined as follows. If *x* is the distance between the upper bound of the tolerance zone and the maximum, then peak duration is the difference between the times at which 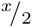 is crossed first (during attainment of the peak) and it is crossed second (during the descent from the peak). Like peak time, peak duration was also normalized by response time. This metric was adapted from “peak duration” defined by Ahmet et al. in their analysis of output responses to an input pulse for an open loop network^[20]^.

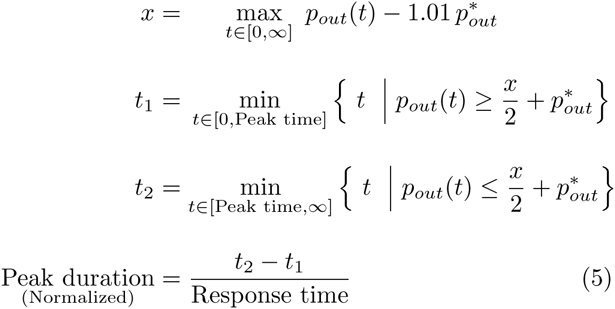

All references henceforth, to overshoot, peak time and peak duration would mean their normalized values.

A perfectly adapting system would have a steady state gain of zero. In order to study the sensitivity of the different motifs to perturbation in different internal parameters (all parameters except input), the calculation of the abovementioned metrics (Equations 1–5) were done with 10000 randomly sampled parameter sets, which gave a distribution of these metrics. These analyses for feedforward and feedback motifs are presented in following sections. The unregulated system is not considered in these sections but some of its properties are shown in Fig. 4C, for comparison.

**Fig. 4:**
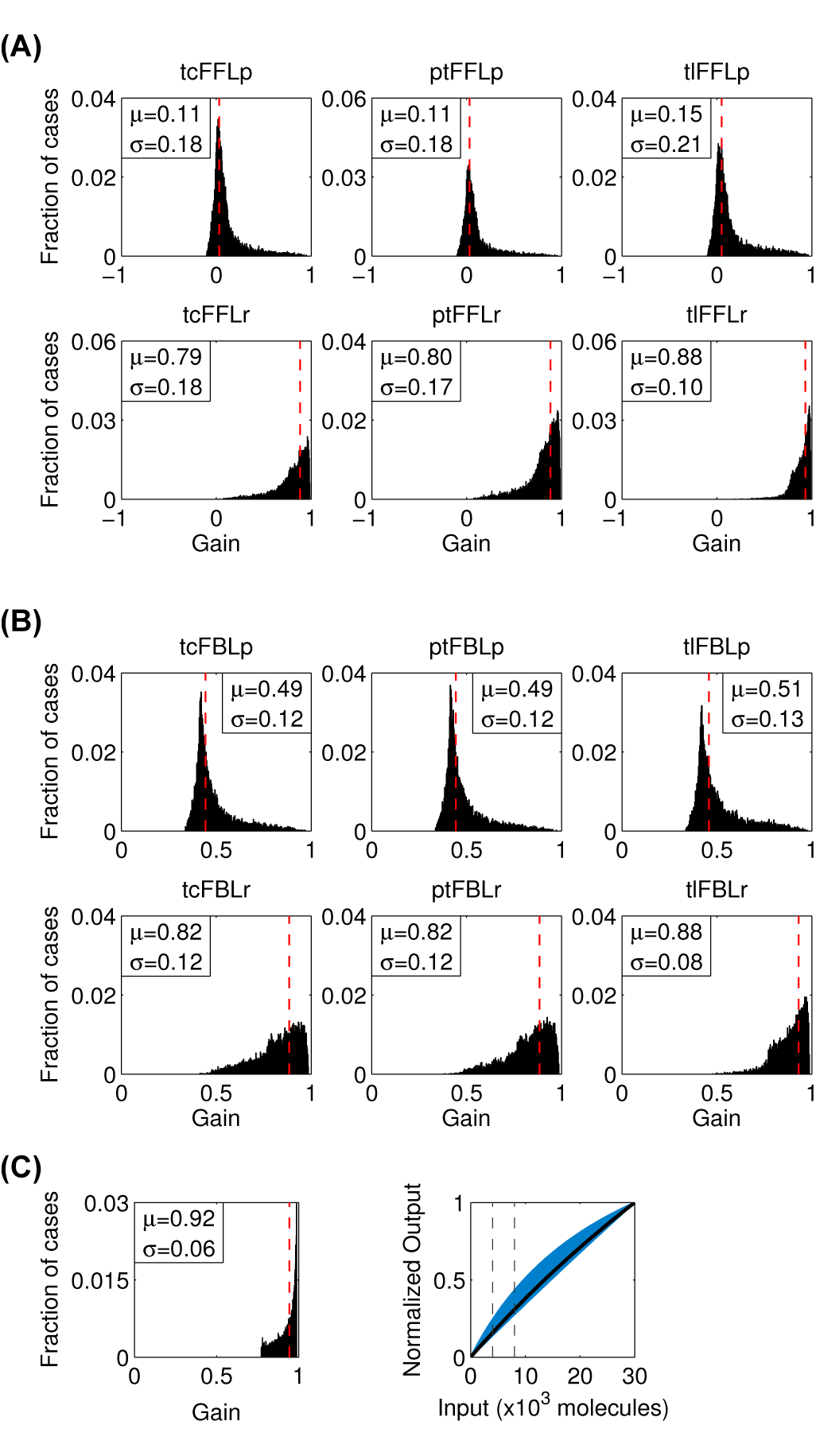
Steady state gain distributions for (**A**) feedforward motifs and (**B**) feedback motifs. Distribution of steady state gain to input step change from 4000 to 8000 molecules, with randomly sampled parameter sets. The dashed vertical lines denote the gain values corresponding to the default parameter set. *μ* and σ denote mean and standard deviation of the distribution, respectively. Subfigure (**C**) shows steady state gain distribution and input-output characteristics of the unregulated motif (refer Fig. 5).

**Fig. 5:**
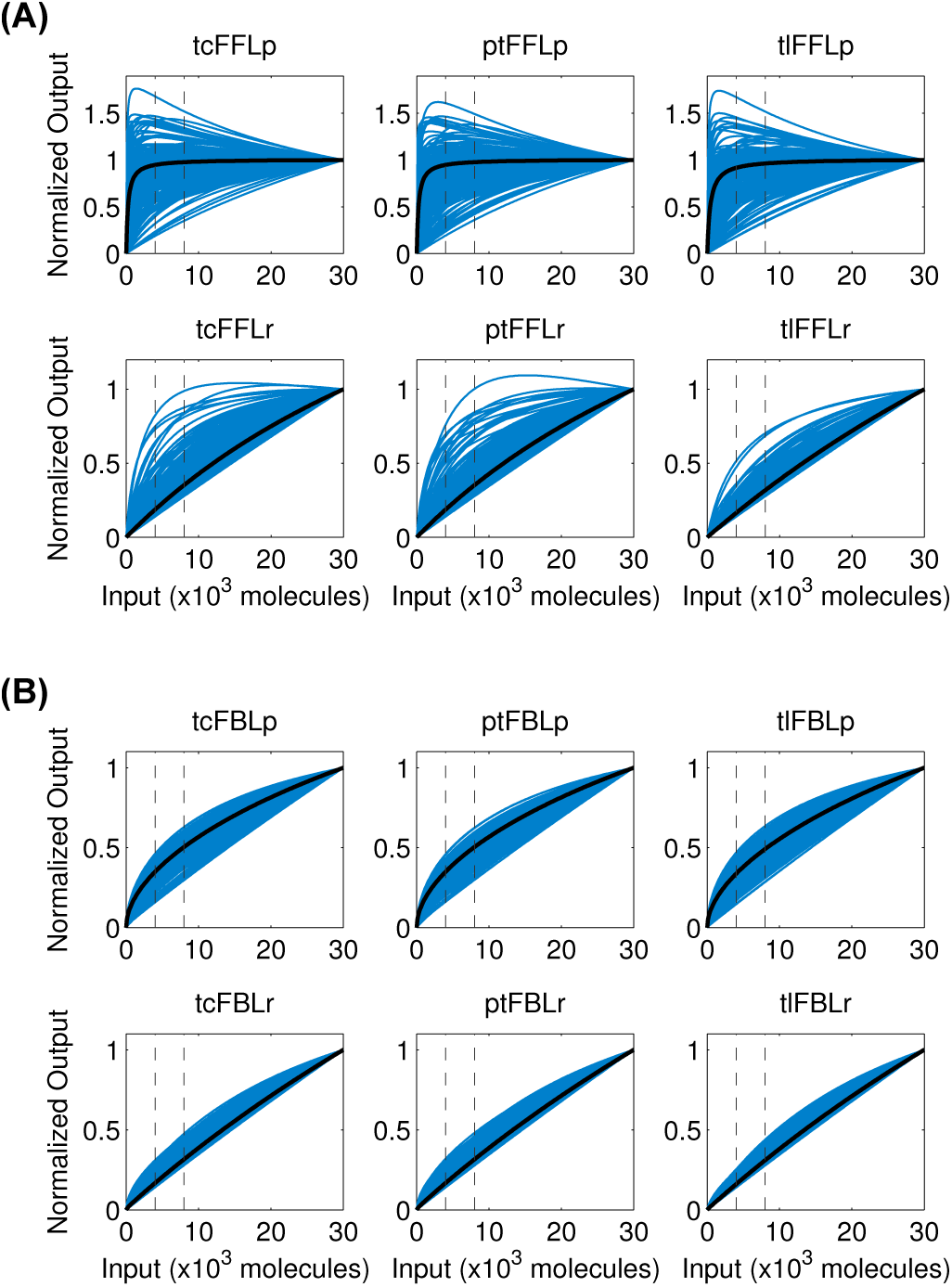
Input-output characteristics for 200 randomly selected cases of (**A**) feedforward and (**B**) feedback motifs. The steady state values at each value of input, are normalized to the value at the maximum value of input (30 × 10^3^). Black line represents the characteristics of the default parameter set. Dashed vertical lines (from left to right) denote input corresponding to 4000 and 8000 molecules respectively.

### 2.2 Mathematical models for the motifs

The mathematical models are described by coupled ordinary differential equations (ODE) with *r*_*out*_, *r*_*ctrl*_, *p*_*out*_ and *p*_*ctrl*_ as variables. The input was considered to be a constant. The ODEs for different motifs are of this general form:

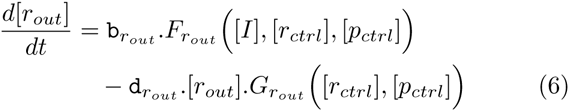

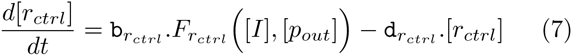

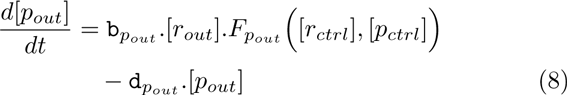

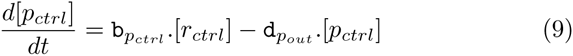

Here, b_*r*_*out*__ and d_*r*_*out*__ denote formation and degradation rate constants for *r*_*out*_ (similarly for others). The functions, *F*_*r*_*out*__, *G*_*r*_*out*__, *F*_*p*_*out*__ and *F*_*r*_*ctrl*__, for different motifs are described in Tab. S1.

### 2.3 Parameter sampling and simulation

A default parameter set (Tab. S2) was chosen based on data obtained from literature. 10000 parameter sets were randomly sampled between an interval around the default value (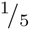 to 5 times the default value) to generate sets with different combination of parameter values. The sampling was done in the log transformed range to avoid bias towards larger samples. These 10000 sets were generated once using the sampling technique described above, and the same sets were used for simulations of different motif structures and regulation mechanisms. The model was simulated using Matlab (ver 7.6.0) ode15s solver for temporal dynamics and steady state values were obtained analytically. In step-up simulations, the initial values of the variables were set to the steady state values at a low input value (corresponding to 4000 molecules/cell). The response to setting the input to 8000 molecule/cell was calculated by numerically integrating the ODEs described in the equations 6–9.

Simulations and calculations were also done with the default degradation rate constant of both the proteins set to a lower (0.1× the original value) or a higher (10×, 100× and 1000×d_*p*_) value and the 10000 parameters sets sampled around each of these new baseline values.

## 3 Results and Discussion

### Protein mediated motifs show higher adaptation in response to input compared to RNA mediated motifs

We used analytical expressions (Supplementary Section 1.3) for steady state to calculate the values of gain (Eqn. 1) for different modes of motifs, for 10000 parameter sets with each parameter varying from 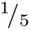 to 5 times the default value. As shown in Fig. 4A, protein mediated motifs exhibited higher level of adaptation (indicated by low gain) compared to the RNA mediated motifs, which had a mild regulatory effect. This behaviour was consistent when the parameters were varied in a broader range (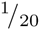 to 20 times the default values; Fig. S1). However, protein mediated FFLs but not FBLs showed perfect or near-perfect adaptation which set the two motif structures apart. Moreover, the gain distribution for FFLs was broader than that of the FBLs with corresponding mode of regulation.

We also tested this response for a range of protein abundances. The gain distributions for protein mediated FFLs were narrower with a strong peak at zero for higher protein abundances (0.1×d_*p*_) while the corresponding FBL motifs were unchanged from the 1× case (Fig. S7-S8). With decreasing protein abundance the gain distribution of all protein mediated motifs broadened (10×d_*p*_, 100×d_*p*_) and finally shifted towards 1 (1000×d_*p*_). The gain of RNA mediated FFLs were unaffected whereas the gain distribution of RNA mediated FFLs broadened with lower mean values for 10 × d_*p*_ and 100 × d_*p*_ cases. As expected at very low protein abundance (1000×d_*p*_) all the regulatory mechanisms had a very mild effect and the response tends to the unregulated response.

To see the effect of magnitude of input on gain, the output was plotted as a function of input (Fig. 5A, Fig. S2A). These plots (input-output or i/o characteristics) reveal that the protein mediated FFLs show adaptability for a wide range of inputs. The shape of i/o characteristics for these motifs have the shape of a saturating function and the steady state value of the output saturates at a low value of input for most parameter combinations. A few cases of tcFFLr and ptFFLr also showed the saturating behaviour. The i/o characteristics, especially in case of the protein mediated FFLs were non-monotonic for some parameter sets – the output initially increased with the input and then decreased, though the descent was not as steep as the ascent. This suggests that the gain and sensitivity for FFLs depend on the the choice of initial and final input values. Contrastingly, FBLs (Fig. 5B) showed monotonically increasing i/o characteristics which did not saturate for the range of the input considered for this analysis (0 to 3×10^4^ molecules); they remained monotonic even with parameter variation in the broader range (Fig. S2B). With lower protein abundance, the non-monotonicity and saturation behaviour of the i/o characteristics of protein mediated FFLs diminished whereas there was no significantly apparent change in the i/o characteristics of FBLs (Fig. S9-S10).

### Protein mediated motifs exhibit high peaking tendency, irrespective of the motif structure and mode of regulation

A motif is defined to exhibit peak if it has a non-zero overshoot (Eqn. 3). Both feedforward and feedback motifs show peaks and protein mediated motifs exhibit a high propensity to give peaks, as reflected by the total number of peaks observed for the 10000 parameter combinations (Fig. 6). The RNA mediated motifs had very low peaking tendency. Overall, peaking propensity only depended on the nature of the controller. However, the peaking tendencies of protein mediated motifs decreased while that of the RNA mediated motifs increased with decreasing protein abundance (Fig. S11-S12). With increased abundance, the effect is even more pronounced with none of the RNA mediated motifs exhibiting peaks.

**Fig. 6:**
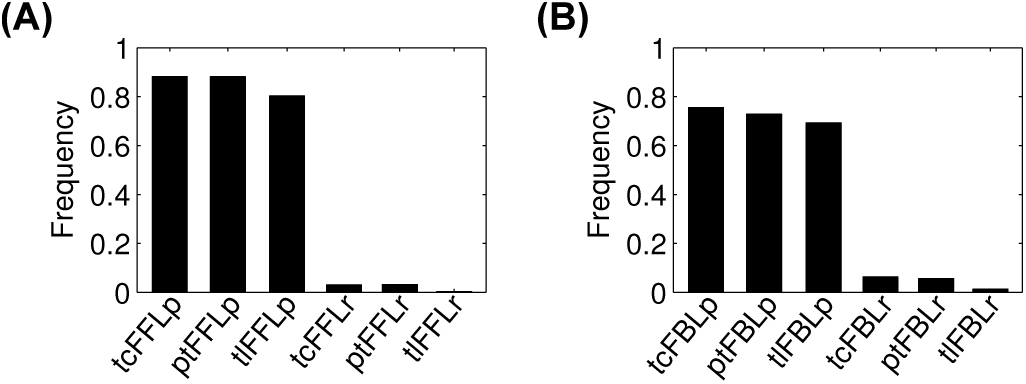
Peaking tendencies of (**A**) feedforward and (**B**) feedback motifs. Y-axis denotes fraction of parameter sets among the 10000 that generate peaks.

To understand the characteristics of these peaks, their properties i.e. overshoot, peak time (Eqn. 4) and peak duration (Eqn. 5) were measured. As mentioned previously, peak time and peak duration were normalized to response time (distributions shown in Fig S3) and overshoot was normalized to the final steady state. Since tlFFLr and tlFBLr showed a very low peaking tendency, they are excluded from further analyses (their properties are shown in Fig. S4).

All RNA mediated motifs, both FFL and FBL, showed a very low mean overshoot which was close to the cut-off limit i.e. 0.01 (Fig. 7). The protein mediated motifs had a much broader distribution and higher mean overshoot. Though the overshoot distribution for both protein mediated FFLs and FBLs was skewed towards zero, the former was broader with higher mean values of overshoot.

**Fig. 7:**
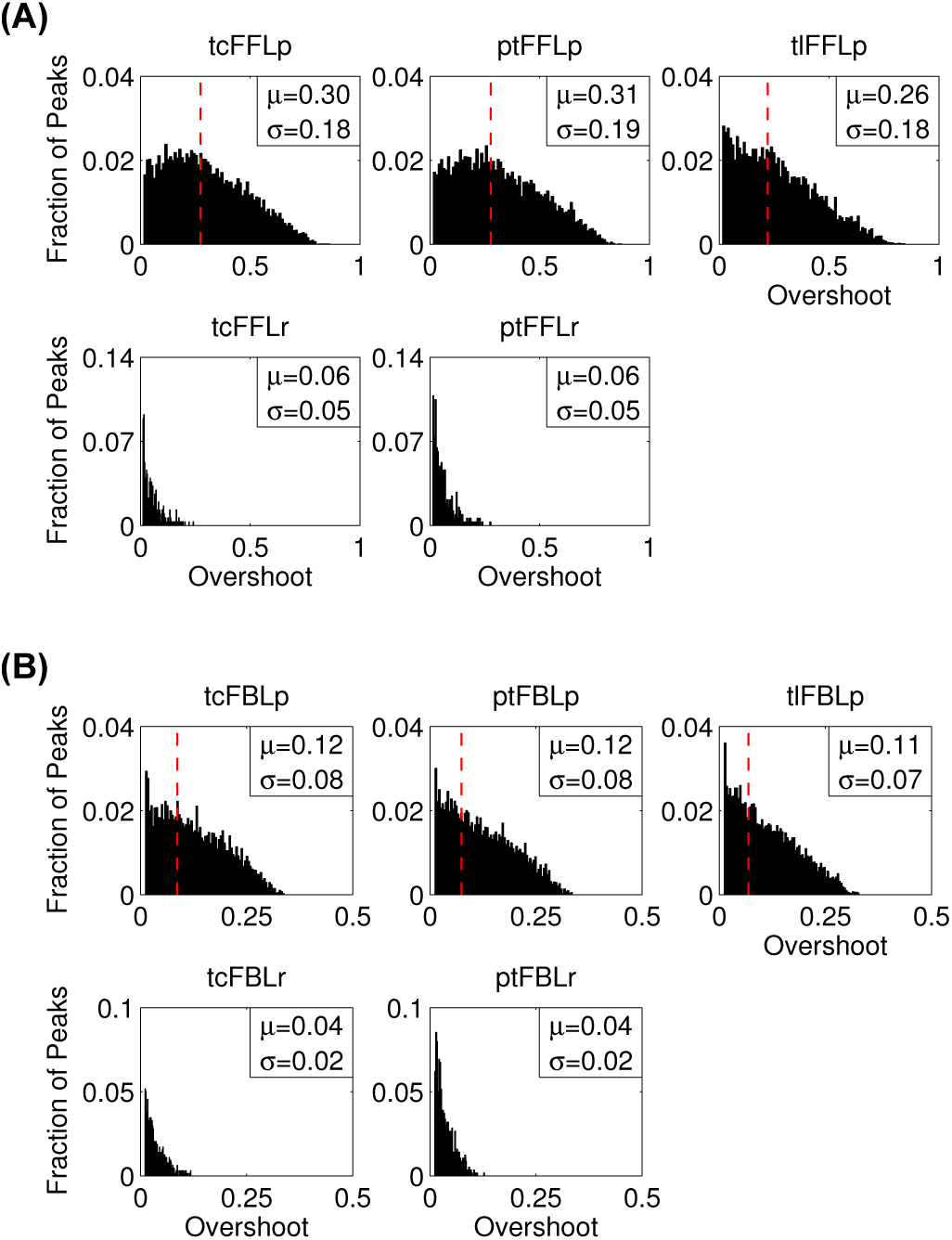
Distributions of overshoot for the parameter sets that generate peaks in (**A**) feedforward and (**B**) feedback motifs. The dashed vertical lines denote values corresponding to the default parameter set if it exhibits peak. Y-axis denotes the fraction of cases among the total number of instances that produce peaks. *μ* and σ denote mean and standard deviation of the distribution, respectively.

The peak time distributions (Fig. 8), however, followed opposite trends between protein mediated FFLs and FBLs. The distributions were narrower in case of protein mediated FFLs with mean peak time at ~0.18 while that for corresponding FBLs were much broader with mean peak time at ~0.4. RNA mediated motifs, both FFL and FBL, showed higher mean peak time than the protein mediated motifs. However, the RNA mediated FBLs had a higher mean peak time and a narrower distribution compared to corresponding FFLs. It can be noted that peak time and overshoot depend on the nature of controller and the motif structure but not to a significant extent on the mode of regulation.

**Fig. 8:**
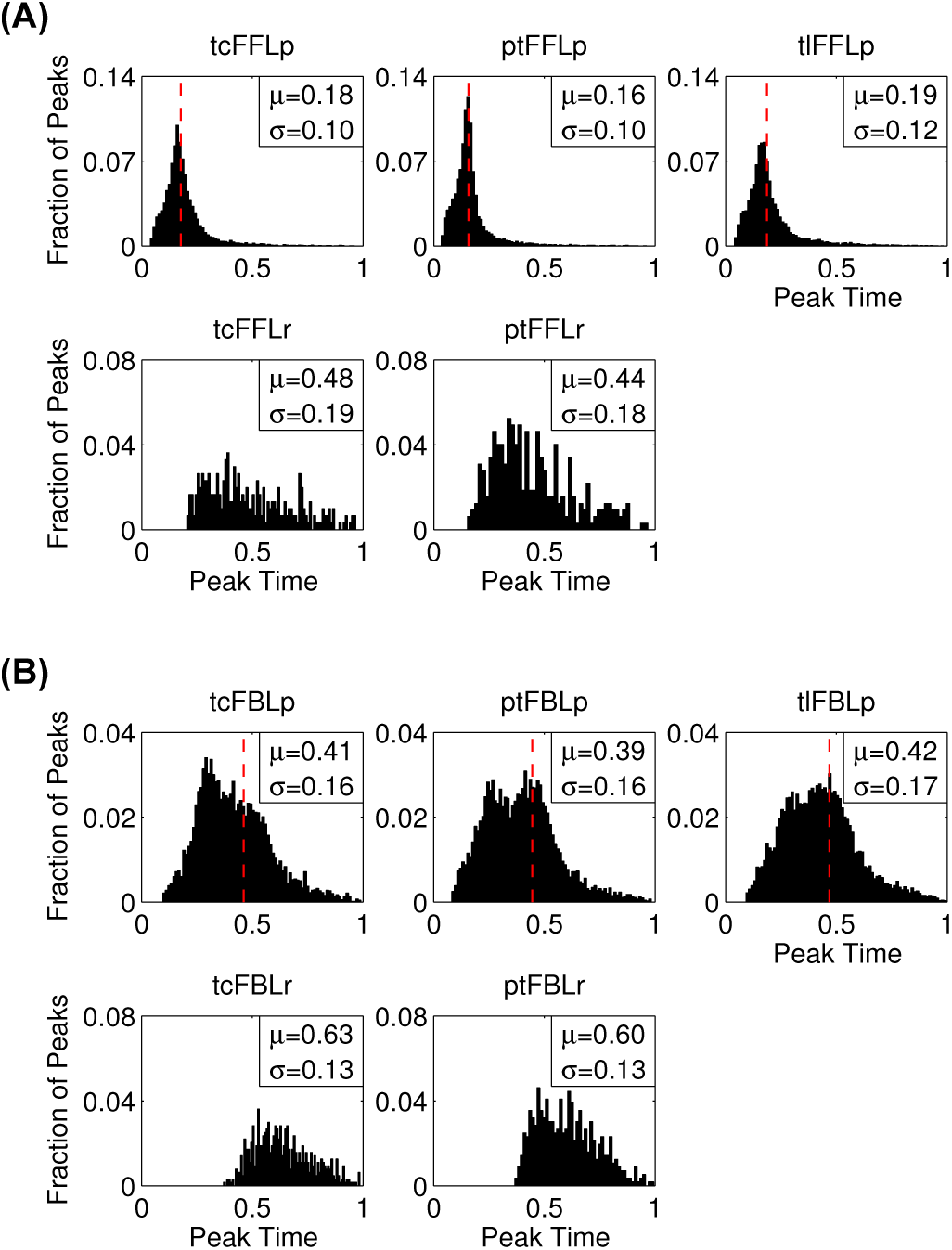
Distributions of peak time for the parameter sets that generate peaks in (**A**) feedforward and (**B**) feedback motifs. The dashed vertical lines denote values corresponding to the default parameter set if it exhibits peak. Y-axis denotes the fraction of cases among the total number of instances that produce peaks. *μ* and σ denote mean and standard deviation of the distribution, respectively.

Overshoot distributions for all motifs moved closer to zero with decreasing protein abundance (Fig. S13-S14) while the peak time distribution for protein mediated FFLs broadened (Fig S15-S16).

The distributions for the peak duration (Fig. 9B) for the protein mediated feedbacks, exhibited apparent bi-modality for the range of parameter variation at 1x protein degradation rates. All of them showed peaks close to ~0.3 and ~0.45, but the ratio of first peak to second peak reduced from tcFBLp, ptFBLp to tlFBLp. For all these motifs, the value of peak duration corresponding to the default parameter set overlapped with the second peak. However, this bimodality seems to be dependent on protein abundances as it could not be seen for the 0.1x, 100x and 1000x cases (Fig. S18). In contrast, the distributions of peak duration for feedforward motifs showed one sharp peak near 0.35. Moreover, the nature of the regulator (Fig. 9A) and protein abundances (Fig. S17) seemed to have no significant effect on the peak duration distribution of FFLs.

**Fig. 9:**
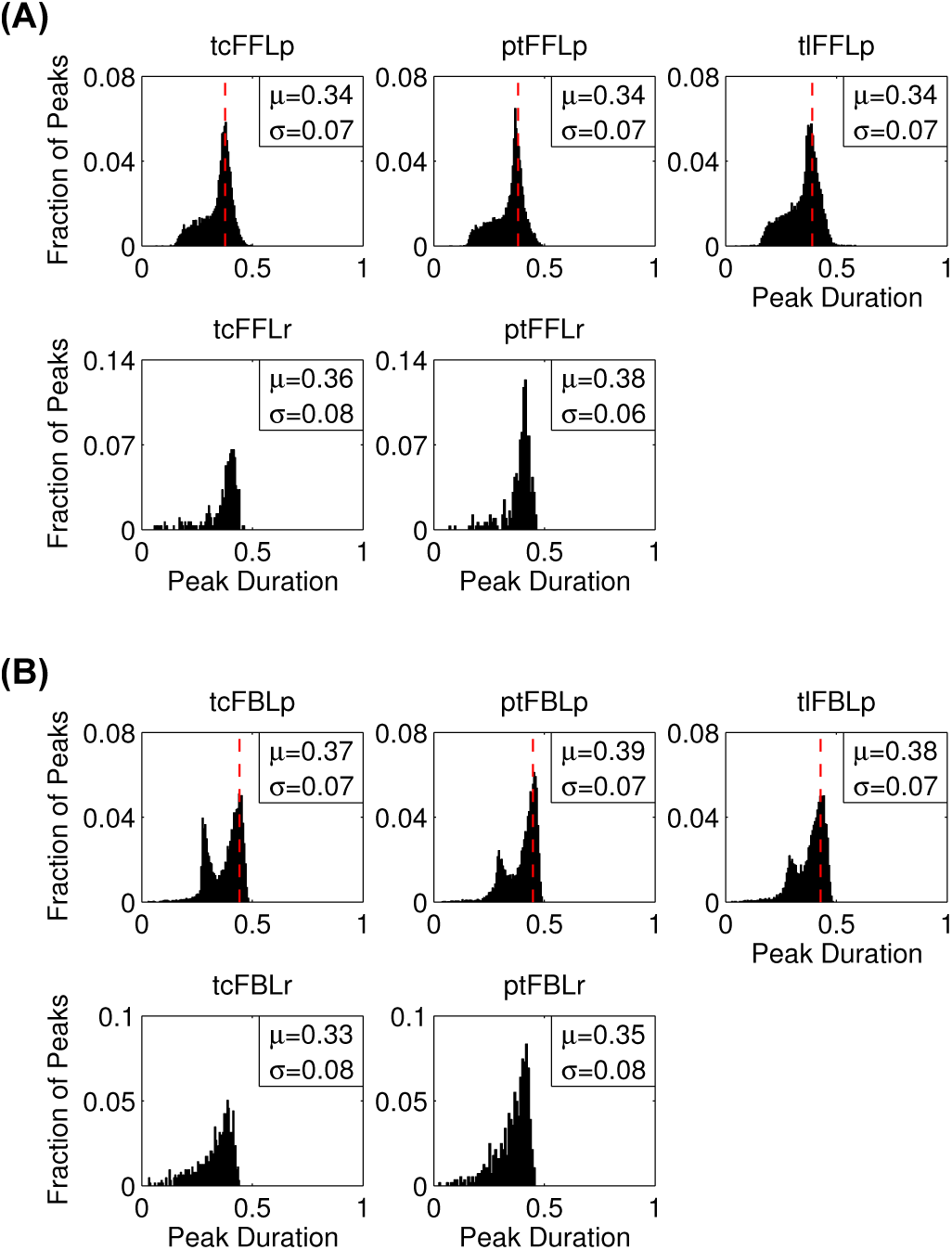
Distributions of peak duration for the parameter sets that generate peaks in (**A**) feedforward and (**B**) feedback motifs. The dashed vertical lines denote values corresponding to the default parameter set if it exhibits peak. Y-axis denotes the fraction of cases among the total number of instances that produce peaks. *μ* and σ denote mean and standard deviation of the distribution, respectively.

### Overshoot decreases whereas peak duration concavely varies with peak time

In order to have a better understanding of the peak dynamics the relationship between overshoot, peak duration and peak time was studied using scatter plots. For both feedforward and feedback motifs, the overshoot apparently decreases exponentially with peak time, though the slope is steeper in case of the FFLs (Fig. 10). This is clearly evident in case of protein mediated motifs but not so much in case of RNA mediated motifs because they exhibit a very low overshoot.

**Fig. 10:**
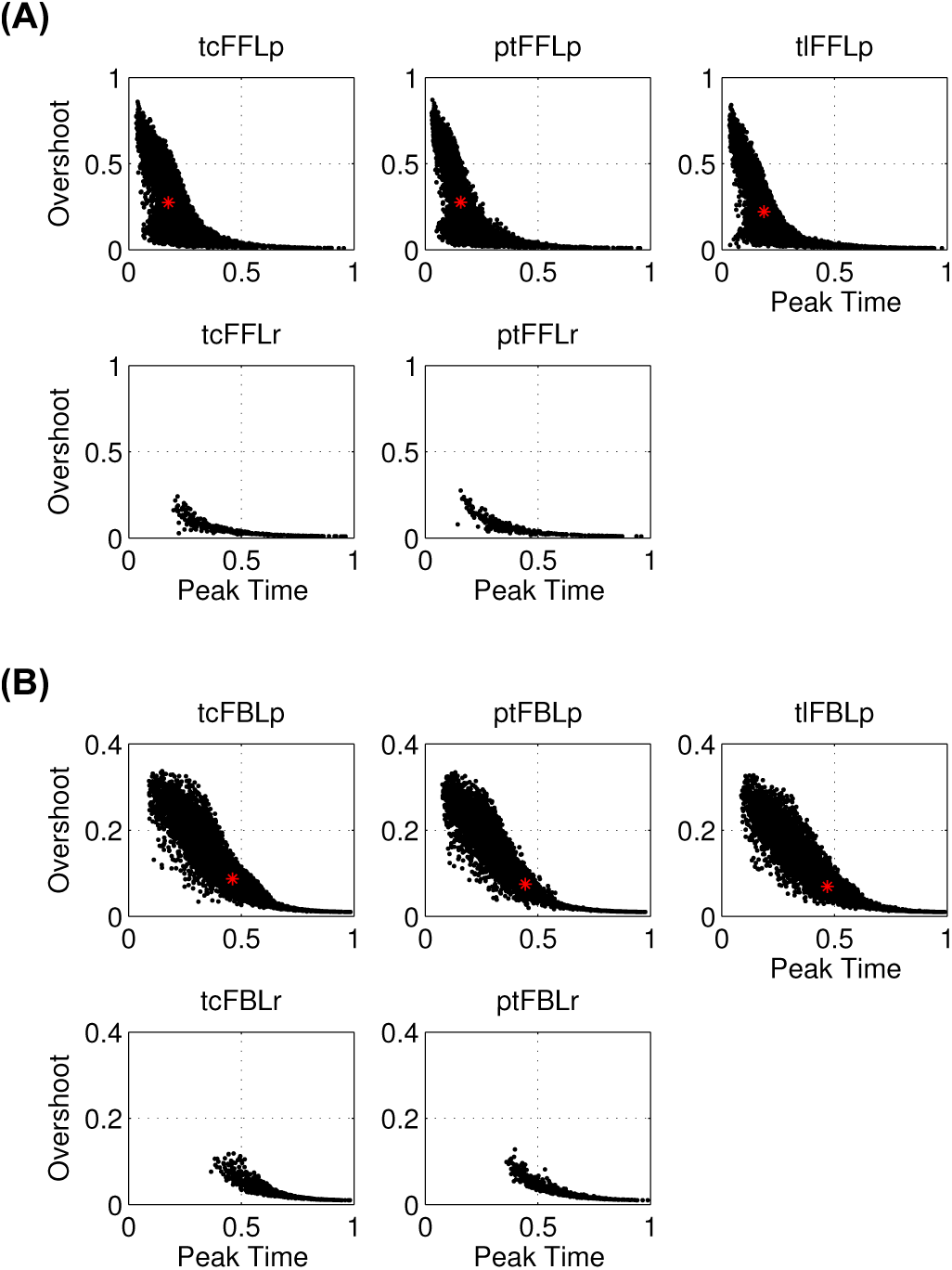
Scatter plots between peak time and overshoot for (**A**) feedforward and (**B**) feedback motifs. Asterisks denote the values corresponding to the default parameter set if it exhibits peak

The scatter plots between peak duration and peak time (Fig. 11) showed a very interesting observation that peak duration initially increases with peak time and then decreases. In other words peak duration seems to be a concave function of peak time. This relationship was observed in both protein mediated feedback and feedforward motifs and was independent of the mode of regulation. This could also be seen in case of RNA mediated FFL, although not as strikingly as the protein mediated motifs. However, because RNA mediated FFLs show delayed peaks, the initial increase of peak duration with peak time could not be observed.

**Fig. 11:**
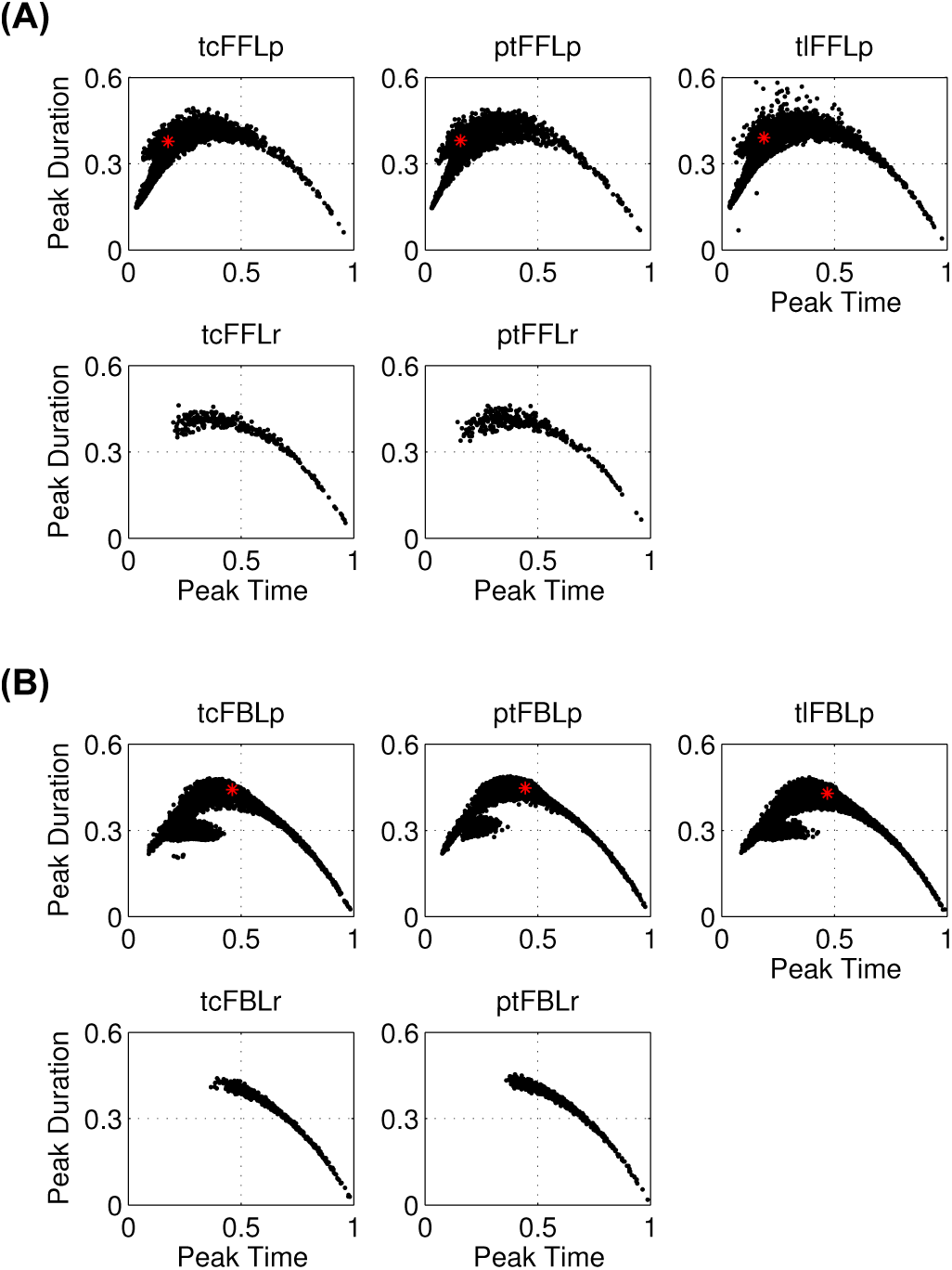
Scatter plots between peak time and peak duration for (**A**) feedforward and (**B**) feedback motifs. Asterisks denote the values corresponding to the default parameter set if it exhibits peak

Both these relationships were unaffected by protein abundances (Fig. S19-S22).

## 4 Conclusion

In this study, we have used a simple and widely applied model for gene expression to enable a direct comparison between the incoherent-feedforward and negative feedback motifs in which the control is implemented at different levels of gene expression. To our knowledge, such a study using a common model has not been reported. The use of a common model with common parameter sets allows a direct comparison between the steady state and the dynamic properties of feedback and feedforward motifs.

Our simulations reveal some previously reported features such as adaptability, that are common between different modes of regulation. They also reveal features that are common to diverse kinds of motifs and depend on the level at which the control is implemented and inter-metric relationships that are independent of motif structure and mode of regulation.

It was observed that while some properties depended on the type of the motif, others were dependent on the mode of regulation. Overall, it is apparent that all protein mediated feedforward motifs can show adaptation whereas feedback motifs just tone down the output response. Though negative integral feedback has been shown to exhibit adaptation ^[14, 21]^, the analytical expressions for simple negative feedbacks considered in this study show that these motifs do not exhibit adaptation. It is interesting to note that while bacterial chemotaxis employs the negative integral feedback, eukaryotic cells such as *Dictyostelium* use the incoherent feedforward loop^[22, 23]^. However, adaptability is not always desirable; in many situations the system is required to respond, but in a controlled manner. Protein mediated feedforward motifs show perfect adaptation at different input ranges because input-output characteristics saturates at a very low value of input. However, since the saturation point is so small, any small stochastic increase from zero input will lead to a sharp response which might be a trade-off.

RNA mediated control has a very mild effect and perhaps just serves as a mechanism to fine tune the gene expression^[24, 25, 26]^. The stronger response mediated by regulatory proteins compared to RNAs is possibly because the regulatory signal gets amplified due to translation. This is also supported by the observation that increasing the protein degradation rates, thereby reducing their abundances, reduces their regulatory effect. At very high protein degradation rates, the system effectively behaves like an unregulated system. miRNAs are known to control regulation by both mRNA degradation and translational inhibition^[27]^. The translational mode of regulation would be effective when the mRNA concentration is less and the protein synthesis rate per mRNA is high. In case of cells like neurons where the nucleus is far from some cellular compartments like the axon-termini and dendritic spines, translational inhibition, which happens locally, would be faster than transcriptional control, which happens at the soma. Even though post-transcriptional regulation might repress the expression as fast as (or faster than) translational regulation, de-repression would still be slow because mRNA re-accumulation would require its transport from the soma. It has been shown experimentally, that mRNAs that are translationally repressed by miRNAs in the synaptic compartments of neurons, are de-repressed upon signalling via the NMDA-receptor^[28]^. Overall, it can be concluded that adaptation and i/o characteristics are dependent on the motif structure and the nature of the controller.

Ability to generate pulses is a useful feature in certain conditions where the response should not persist at a high level for a long time. Examples of such conditions include heat shock and other kinds of acute stresses^[29]^. Both feedforward and feedback motifs can generate pulses of which the former can also adapt back to the original state, as discussed previously. Overshoot reduces with peak time, suggesting that faster the peak is attained, higher will be its impact. The peak duration initially increases with peak time and attains a maximum before decreasing, however, the overshoot is relatively low at these values of peak time. Therefore a suboptimal peak duration may be better for an effective response. In protein mediated feedback motifs, the peak duration shows apparent bimodality for certain range of parameters. It may be possible to switch from long lasting peaks to sharp peaks by tweaking the parameters using an extrinsic agent.

It can be concluded that while dynamic properties such as peak time and peak duration are strongly dependent on the motif structure as well as the mode of regulation, peaking ability itself seems to be dependent only on the nature of regulator. Moreover, the relationships between peak duration, overshoot and peak time seem to follow a paradigm that is universal to all the considered motifs.

## Acknowledgements

Authors acknowledge funding from the Council of Scientific and Industrial Research (CSIR), India (BRI; Senior Research Fellowship), CSIR-Institute of Genomics and Integrative Biology (BP; project BSC0123) and Department of Biotechnology (DBT), India (KVV; project ref. BT/EB/PAN IIT/2012, Dt. 08/12/2014).

## References

[1] S. Gokhale, D. Nyayanit, andC. Gadgil, “A systems view of the protein expression process”, Systems and Synthetic Biology, vol. 5, no. 3-4, pp. 139–150, 2011.

[2] Q. Liu andZ. Paroo, “Biochemical principles of small RNA pathways”, Annual Review of Biochemistry, vol. 79, no. 1, pp. 295–319, 2010. PMID: 20205586.

[3] J. L. Rinn and H. Y. Chang, “Genome regulation by long noncoding RNAs”, Annual Review of Biochemistry, vol. 81, no. 1, pp. 145–166, 2012. PMID: 22663078.

[4] S. S. Shen-Orr, R. Milo, S. Mangan, and U. Alon, “Network motifs in the transcriptional regulation network of Escherichia coli”, Nat Genet, vol. 31, pp. 64–68, May 2002.

[5] U. Alon, “Network motifs: theory and experimental approaches”, Nat Rev Genet, vol. 8, pp. 450–461, Jun 2007.

[6] J. J. Tyson and B. Novák, “Functional motifs in biochemical reaction networks”, Annual Review of Physical Chemistry, vol. 61, no. 1, pp. 219–240, 2010.

[7] S. Mangan and U. Alon, “Structure and function of the feed-forward loop network motif”, Proceedings of the National Academy of Sciences, vol. 100, no. 21, pp. 11980– 11985, 2003.

[8] E. Sontag, “Remarks on feedforward circuits, adaptation, and pulse memory”, Systems Biology, IET, vol. 4, pp. 39–51, Jan 2010.

[9] L. Goentoro, O. Shoval, M. W. Kirschner, and U. Alon, “The incoherent feedforward loop can provide fold-change detection in gene regulation”, Molecular Cell, vol. 36, pp. 894–899, Dec 2009.

[10] P. Wang, J. Lü, and M. J. Ogorzalek, “Global relative parameter sensitivities of the feed-forward loops in genetic networks”, Neurocomputing, vol. 78, no. 1, pp. 155 – 165, 2012. Selected papers from the 8th International Symposium on Neural Networks (ISNN 2011).

[11] M. Duk, M. Samsonova, and A. Samsonov, “Dynamics of miRNA driven feed-forward loop depends upon miRNA action mechanisms”, BMC Genomics, vol. 15, no. Suppl 12, p. S9, 2014.

[12] W. Ma, A. Trusina, H. El-Samad, W. A. Lim, and C. Tang, “Defining network topologies that can achieve biochemical adaptation”, Cell, vol. 138, no. 4, pp. 760– 773, 2009.

[13] S. Iadevaia, L. K. Nakhleh, R. Azencott, and P. T. Ram, “Mapping network motif tunability and robustness in the design of synthetic signaling circuits”, PLoS ONE, vol. 9, p. e91743, Mar 2014.

[14] T.-M. Yi, Y. Huang, M. I. Simon, and J. Doyle, “Robust perfect adaptation in bacterial through integral feedback control”, Proceedings of the National Academy of Sciences, vol. 97, no. 9, pp. 4649–4653, 2000.

[15] N. Rosenfeld, M. B. Elowitz, and U. Alon, “Negative autoregulation speeds the response times of transcription networks”, Journal of Molecular Biology, vol. 323, no. 5, pp. 785 – 793, 2002.

[16] J. Tsang, J. Zhu, and A. van Oudenaarden, “MicroRNA-mediated feedback and feedforward loops are recurrent network motifs in mammals”, Molecular Cell, vol. 26, no. 5, pp. 753 – 767, 2007.

[17] B. Schwanhäusser, D. Busse, N. Li, G. Dittmar, J. Schuchhardt, J. Wolf, W. Chen, and M. Selbach, “Global quantification of mammalian gene expression control”, Nature, p. 337342, May 2011.

[18] K. Ogata, Modern control engineering. Boston, MA: Prentice-Hall, 2010.

[19] G. Stephanopoulos, Chemical process control: an introduction to theory and practice. Englewood Cliffs, N.J: Prentice-Hall, 1984.

[20] A. Ay, N. Wilner, and N. Yildirim, “Mathematical modeling deciphers the benefits of alternatively-designed conserved activatory and inhibitory gene circuits”, Mol. BioSyst., vol. 11, pp. 2017–2030, 2015.

[21] P. R. Somvanshi, A. K. Patel, S. Bhartiya, and K. V. Venkatesh, “Implementation of integral feedback control in biological systems”, Wiley Interdisciplinary Reviews: Systems Biology and Medicine, vol. 7, no. 5, pp. 301–316, 2015.

[22] C. J. Wang, A. Bergmann, B. Lin, K. Kim, and A. Levchenko, “Diverse sensitivity thresholds in dynamic signaling responses by social amoebae”, Science Signaling, vol. 5, no. 213, pp. ra17–ra17, 2012.

[23] K. Takeda, D. Shao, M. Adler, P. G. Charest, W. F. Loomis, H. Levine, A. Groisman, W.-J. Rappel, and R. A. Firtel, “Incoherent feedforward control governs adaptation of activated ras in a eukaryotic chemotaxis pathway”, Science Signaling, vol. 5, no. 205, pp. ra2–ra2, 2012.

[24] A. S. Flynt and E. C. Lai, “Biological principles of microRNA-mediated regulation: shared themes amid diversity”, Nat Rev Genet, vol. 9, pp. 831–842, Nov 2008.

[25] M. P. Gantier, A. J. Sadler, and B. R.G. Williams, “Fine-tuning of the innate immune response by microR-NAs”, Immunol Cell Biol, vol. 85, pp. 458–462, Jul 2007.

[26] L. Harfouche, F. Z. Haichar, and W. Achouak, “Small regulatory RNAs and the fine-tuning of plant-bacteria interactions”, New Phytologist, vol. 206, no. 1, pp. 98– 106, 2015.

[27] M. A. Valencia-Sanchez, J. Liu, G. J. Hannon, and R. Parker, “Control of translation and mRNA degradation by miRNAs and siRNAs”, Genes & Development, vol. 20, no. 5, pp. 515–524, 2006.

[28] S. Banerjee, P. Neveu, and K. S. Kosik, “A coordinated local translational control point at the synapse involving relief from silencing and {MOV10} degradation”, Neuron, vol. 64, no. 6, pp. 871 – 884, 2009.

[29] N. Yosef and A. Regev, “Impulse Control: Temporal Dynamics in Gene Transcription”, Cell, vol. 144, no. 6, pp. 886 – 896, 2011.

